# Modeling the overproduction of ribosomes when antibacterial drugs act on cells

**DOI:** 10.1101/024703

**Authors:** Arijit Maitra, Ken A. Dill

## Abstract

Bacteria that are subjected to ribosome inhibiting antibiotic drugs show an interesting behavior: Although the drug slows down cell growth, it also paradoxically increases the cell’s concentration of ribosomes. We combine a prior nonlinear model of the energy-biomass balance in undrugged *E. coli* cells (*Maitra and Dill, PNAS 2015*) with Michaelis-Menten binding of drugs that inactivate ribosomes. Predictions are in good agreement with experiments on ribosomal concentrations and synthesis rates vs. drug concentrations and growth rates. The model indicates that added drug drives the cell to overproduce ribosomes keeping roughly constant the level of ribosomes producing ribosomal proteins, an important quantity for cell growth. The model also predicts that ribosomal production rates should increase, then decrease with added drug. This model gives insights into cellular driving forces and suggests new experiments.

## 1. INTRODUCTION

Drugs, such as chloramphenicol, act to reduce bacterial cell growth rates by inhibiting bacterial ribosomes, thereby reducing the cell’s production of proteins. What actions does the cell invoke to counter the drug? On the one hand, there is often a good understanding of how the drug binds at its ribosomal site [1–3] and it is sometimes known how that binding interferes with protein elongation [4–7]. It is also sometimes known how drugs sensitize local networks to evoke adaptive responses [8–11]. On the other hand, there is usually less understanding of what global stresses the drug trigger, how it shifts the balances of energy and biomass, or of what homeostatic condition the cell might be trying to preserve.

There are various approaches to cell-level modeling. One approach models the dynamics of the cell’s networks of biochemical reactions [12–14]. Even in an organism as simple as a bacterium, there are very many interconnected web of reactions, making it complicated to model. Another approach has been Flux-Balance Analysis [15, 16], which gives solutions by linearizing the forces around some given homeostasis point. Here, however, we are interested in how those homeostasis points themselves are shifted by the drug. Homeostasis is a fundamentally nonlinear phenomenon, describing the cell’s return to a stable state after a perturbation. Like the *Le Chatelier Principle* in physics [17], homeostasis describes a process resembling a marble rolling back to the bottom of a well after being pushed, like a well-bottom of an energy function. Here, we treat the nonlinearities and feedbacks that are needed to explore how the homeostasis balance is tipped by the drug. But to do this in a way that can give simple insights, we use a reduced (‘minimalist’) description of the bacterial cell [18]. We use this model to study the response of *E. coli* to chloramphenicol.

Our goal here is a quantitative description of the energy-limited cell in the absence or presence of varying amounts of drug, in terms of the physico-chemical processes of the undrugged cell developed recently [18]. (By energy limited, we mean cells that are growth limited by a sugar source, such as glucose, rather than by amino acids, for example). Our minimal model expresses the dynamical concentrations and fluxes of three internal cell components – ribosomal protein, nonribosomal protein, and internal energy (lumped into a single category we call ATP), as a function of external sugar, such as glucose. We previously found that healthy *E. coli* under good growth conditions (speeds up to one duplication per hour) have achieved an evolutionary balance [18]. On the one hand, the cell invests energy and biomass in increasing its ribosome concentration because that increases the cell’s growth speed. On the other hand, too much energy and biomass devoted to producing ribosomes leads to starving the cell’s ability to take in food and convert it to ATP. In the present paper, we ask how drugging the cell affects its balance of energy and biomass.

## 2. A MINIMAL MODEL OF *E. COLI* IN THE PRESENCE OF DRUGS

We model the energy-limited growth of *E. coli* using three rate equations: for energy (ATP concentration, A) and ribosomal (R*_act_*) and nonribosomal protein concentrations (P) as functions of time *t* [18]:

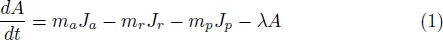

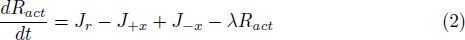

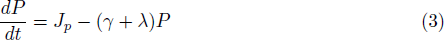

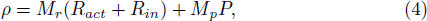

where the fluxes are defined as:

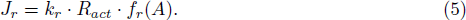

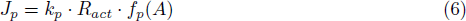

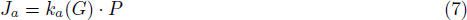

Here, *J_a_* is the rate of glucose conversion for ATP generation, *J_p_* and *J_r_* are the respective rates of synthesis of nonribosomal proteins and ribosomes. *k_r_*, *k_p_* and *k_a_*(*G*) are the respective rate constants for ribosomal biogenesis, protein translation and energy generation. The units of rates and rate constants are ‘mM per hour’ and ‘per hour’ respectively. *m_a_* is the moles of ATP per mole of glucose generated, and, *m_r_* and *m_p_* are the respective moles of ATP consumed per unit mole of ribosome (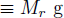 ribosomal proteins) and nonribosomal proteins (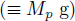) synthesized. Our work is not the first to model the biomass balance in bacteria; see [19–25]. What is new here, we believe, is the coupling between the biomass and energy balance; also see [26].

The functional forms in Eqs. (5), (6), (7) reflect wildtype regulatory mechanisms that coordinate the syntheses of ribosomal and non-ribosomal proteins, which are complex [27, 28] and depend on the cell’s energy status. To capture these dependencies, we adopt the undrugged cell functions [18]:

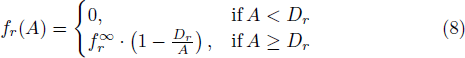

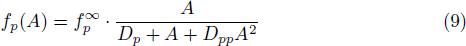

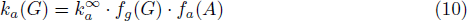

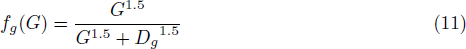

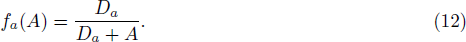

See S.I. for values of biophysical constants, capacities 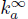, 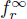, 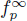, and, parameters *D_r_*, *D_p_*, *D_pp_*, *D_g_* and *D_a_* defining the non-linearities in the respective pathways. In addition, here we consider the effects of drug *X* as shown in Fig. 1. *X* is an antibiotic drug that targets ribosomes. There is a broad class of natural and synthetic bacteriostatic antibiotics of this type, such as chloramphenicol, that target protein synthesis. The present model is intended as a general description of that class of drugs [3]. We assume X permeates *passively* from the extracellular medium into the cytosol through the cell membrane. We assume free drug concentrations outside and inside the cell are equal, a reasonable approximation for *E. coli* based on similar values of drug binding kinetics from *in vivo* and *in vitro* measurements, see Ref. [29, 30].

**Figure 1:**
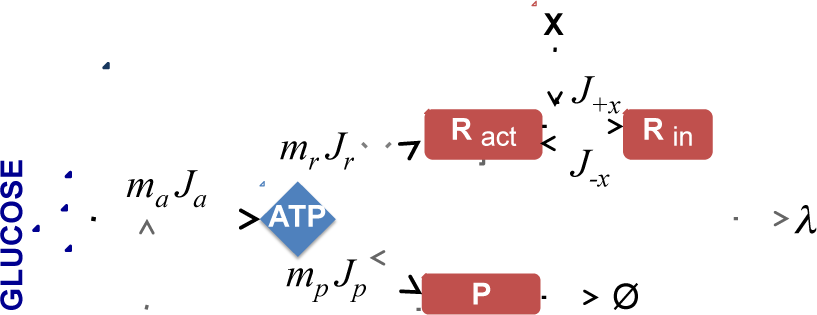
Minimal kinetic model of *E. Coli*. The model expresses the dynamical fluxes (arrows) and concentrations of active ribosomes (R*_act_*), non-ribosomal proteins (P) and a lumped internal energy (ATP). Symbol ⇒ shows a positive feedback mechanism for ribosomal autosynthesis, a key controller of growth behavior. Antibiotic inhibitor molecules are represented by X. X bind reversibly with active ribosomes. While in bound-form, R*_in_*, the ribosomes are inactivated, and they do not translate proteins. P degrades with a rate constant γ. The cell grows exponentially with a specific growth rate of λ.

The binding of X to the ribosomes, which occurs with rate constant *k_+x_*, halts peptide-chain elongation, as represented by the following dynamics:

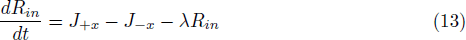

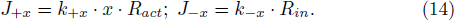

Here, *J*_+*x*_ is the rate at which ribosomes become inactivated due to binding with the drug and *J*_−*x*_ is the rate of unbinding. *R_in_* is the concentration of ribosomes that have been *inactivated* by binding to the drug, and *R_act_*, as noted above, is the intracellular concentration of *active* ribosomes. So, *R_act_* + *R_in_* is the total concentration of ribosomes in the cell. *x* is the extracellular concentration of drugs.

A key quantity in the present model is the fraction of ribosomes that are active *α*(*x*), for a given drug concentration *x*. We assume steady state, so we set 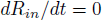 in Eq. (13). We also assume that the rate constant for drug-ribosome unbinding is much faster than dilution 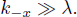. So, we get:

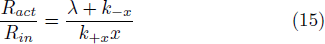

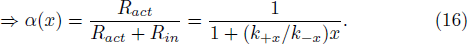

For the equilibrium dissociation constant of chloramphenical, we use 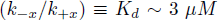 [29]. *α* = 1 represents the situation of no drug. Increasing drug concentration decreases α towards zero.

The fraction of all proteins (by mass) that are active ribosomes is

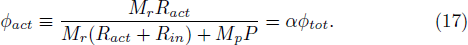

And, the fraction of all proteins that are all ribosomes (active plus inactive) is:

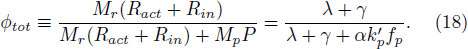

The last equality in Eq. 18 expresses how the ribosomal content of the cell depends on its growth rate λ and other properties. And, then the fraction of active ribosomes devoted to translating ribosomal proteins is:

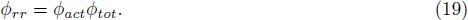

Further, the rate of ribosome synthesis, 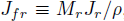, in units of g of ribosomal protein per g of total protein per hour can be computed as (see S.I.):

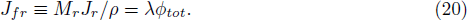

Under growth conditions in the absence of drugs, *α* = 1, and 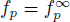 [18] the rate of ribosome synthesis is given as, using Eq. (18) and Eq. (20):

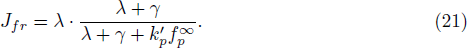

## 3. THE DRUGGED CELL OVERPRODUCES TOTAL RIBOSOMES TO MAINTAIN SUFFICIENTLY MANY ACTIVE RIBOSOMES

Here we describe the model predictions. We solve ODEs (1)-(14) under steady-state conditions for different concentrations of glucose and antibiotic drug. Fig. 2A shows that the model predicts Monod-like behavior [31] of growth rate vs. glucose concentration under different drug concentrations. As expected, the model predicts that increasing drug leads to diminishing maximum-growth rates.

**Figure 2:**
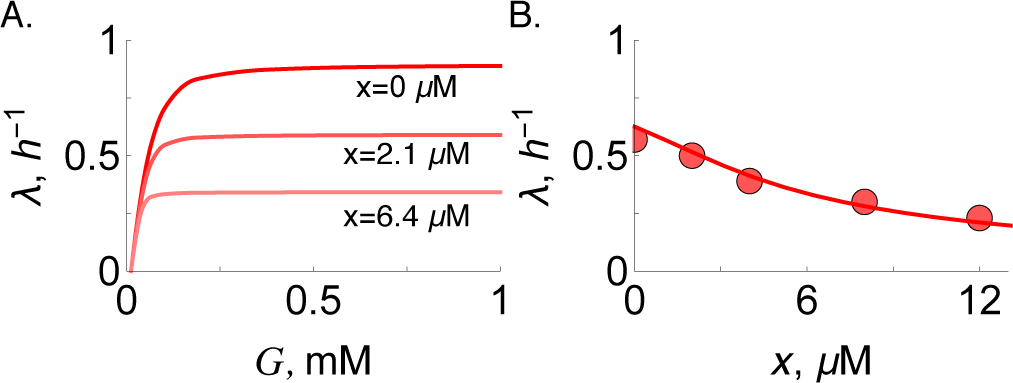
*E. coli* physiological correlations. (A) Growth rate vs. extracellular glucose concentration from simulation for antibiotic (chloramphenicol) concentrations of 0, 2.1 and 6.4 *μ*M. (B) Dependence of growth rate, λ, on antibiotic concentration, *x*. Line is the numerical solution of the ODE model, with *G* = 0.08 mM, red. Red solid circles are experimental data [10, 32] of *E*. *coli* grown on glucose+M63.

Fig. 3 shows that the model is consistent with experiments indicating how added drug stimulates total ribosome production even as it reduces the cell’s growth rate [10, 33]. The black line shows that, for undrugged cells, ribosomes become upshifted relative to other protein biomass with increasing cellular growth rate. The red line and data points show that the added drug does two things: it increases the ribosomal fraction while simultaneously reducing the growth rate.

Our result reduces to the linear model of Scott et al [10] in the limit of zero degradation. To see this, note that (see SI),

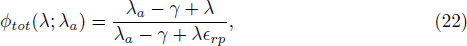

where 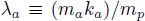 is a measure of specific rate of energy generation and 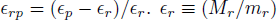 and 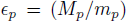 are constants denoting the respective gram-weights of ribosomal and nonribosomal proteins synthesized per mole of ATP. Setting 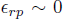 and γ = 0 gives 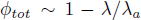, which is just the linear relationship of Scott et al. [10].

**Figure 3:**
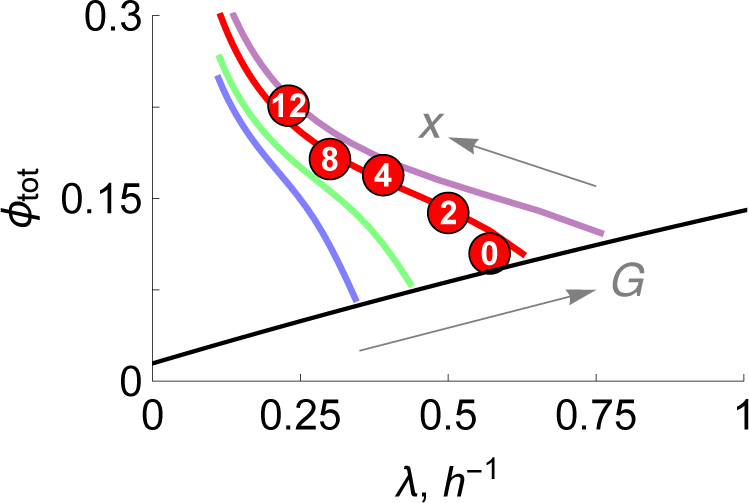
*E. coli* ribosomal protein fraction vs growth rates. Numerical solutions of the ODE model, with increasing glucose concentrations in mM: *G* = 0.04 (blue), 0.05 (green), 0.08 (red), 0.125 (purple), and, antibiotic concentrations *x* = 0 → 25 *μ*M shown by arrows. Circles are experimental data [10, 32] of *E. coli* grown on glucose+M63 at different dosage of chloramphenicol marked in *μ*M. Black line is theory, Eq. (18) with 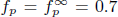, 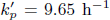, 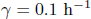, and *α* = 1, absence of drugs.

What is the cell ‘trying to achieve’ under the burden of the drug? As noted above, the effect of the drug is to decrease substantially the fraction of useful ribosomes; see the quantity *α* in Fig. 4. However, Fig. 4 also shows that there is remarkable relative constancy in two other quantities, *ϕ_tot_* and *ϕ_rr_*, independent of the concentration of drug. Ribosomes make either ribosomal or non-ribosomal proteins. *ϕ_rr_* is the fraction of active ribosomes that are producing other ribosomes (see the double arrow in Fig. 1; see also Fig. S2-S4. And, *ϕ_tot_* is the fraction of all proteins that are ribosomal. The constancy of these quantities suggests that the cell senses and regulates how many of its proteins are ribosomes, or how many are ribosomes producing other ribosomes. Such processes may be mediated by ppGpp, the molecule that provides stringent control of ribogenesis in the presence of antibiotic stress [27, 34]. To our knowledge, *ϕ_rr_* has not been measured experimentally.

**Figure 4:**
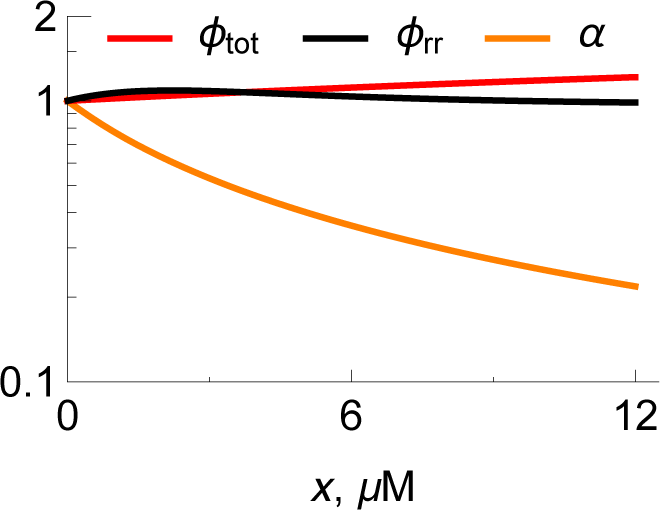
Effect of ribosomal inhibitors on cellular homeostasis. Lines are scaled numerical solutions of the ODE model, with *G* = 0.08 mM. (Orange) *Active* ribosomes, 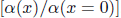, as a function of drug concentration *x*. (Red) *Total* ribosomes, 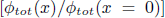. (Black) The fraction of *active ribosomes that are producing ribosomal proteins*, 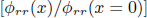; Eq. (19). Also see Fig. S2-S4.

## 4. THE DRUG SHIFTS THE PRODUCTION RATE OF RIBOSOMES. BUT, IT HAS LITTLE EFFECT ON ENERGY FLOW FROM GLUCOSE TO ATP

In this section, we get further insights from looking at two additional properties of the model. First, in Fig. 5, we go beyond *concentrations* of ribosomes and consider the *rate of production* of ribosomes, 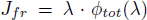 [Eq. (20)], which we also call *ribosomal flux*. We find (see below) that while high drug concentrations increase the numbers of ribosomes, it also reduces the *rates* of ribosome production. Second, Fig. 6 shows that added drug reduces the growth rate by reducing the catabolic conversion of glucose to ATP. Here are the details.

**Figure 5:**
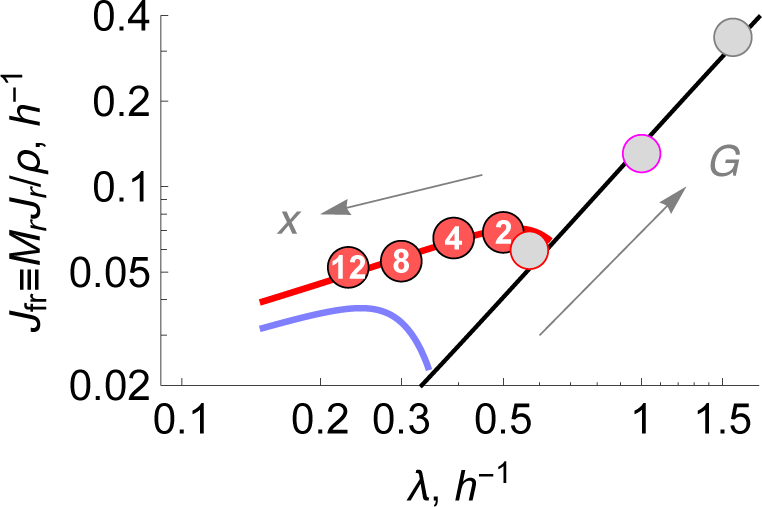
Effect of ribosomal inhibitors on ribosomal activity. The symbols show the rate of ribosomal synthesis 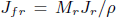 vs. specific growth rate of *E. coli* converted from experimental *ϕ* – *k* data. Chloramphenicol concentrations (*μ*M) marked inside the circles. Nutrients: M63+gluc (red) at T=37 C; Scott et al. [10, 32]. Gray filled circles are experimental data [10, 32] in the absence of drugs. Blue line; ODE model prediction at constant *G* = 0.04 mM and chloromaphenicol varied between *x* = 0 → 15 *μ*M shown by arrow. The black line is theory, Eq. (21), in the absence of drugs with 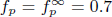, 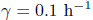, 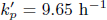. Also see S.I. Fig. S1.

**Figure 6:**
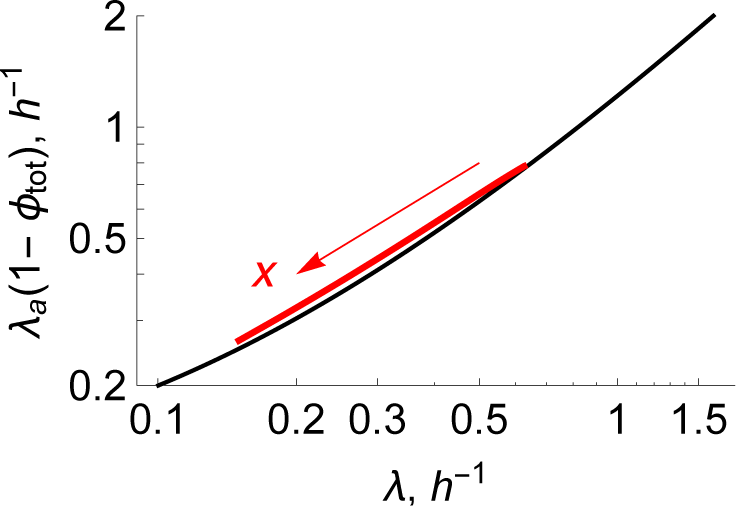
Effect of ribosomal inhibitors on rate of energy metabolism. Predictions of rate of energy metabolism vs. growth rate λ from ODE model under variation of glucose, *G* = 0 – 1 mM and no-drugs (black line). Prediction at *G* = 0.04 mM (red line) under variation of drugs from *x* = 0 → 15 *μ*M shown by the arrow. Increase in drug concentration reduces both rates of growth and energy generation.

First, focus on the black line in Fig. 5. According to the model, under the no-drug condition, the ribosomal production rate should scale as the square of the growth rate, 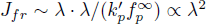, since 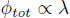. Fig. 5 shows a log-log plot. The black line shows the square-law prediction for undrugged cells. The data points shown in gray lay along this black line, indicating that the model predicts well the ribosomal production rates of undrugged cells growing at different speeds.

Next, focus on the red points in Fig. 5. The datapoints, containing the circled numbers 2-12 *μ*M, show the effects of increasing amounts of drug at fixed nutrients. Following the red line toward the left, which describes increasing drug concentrations, shows how the drug reduces the growth rate while it also reduces the production rate of ribosomes. The experimental datapoints are from ref. [10]; also see SI.

Finally, the blue line on Fig. 5 makes an interesting prediction, for which there are no experiments yet as far as we know. The blue line represents cell growth under low nutrients, 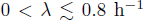. The blue line has curvature. This shows that while ribosomal fluxes are increased by small amounts of added drug, those fluxes are decreased by larger amounts of drug due to the reduction of cell growth at high drug concentrations.

We can draw another inference by comparing the blue and black lines on Fig. 5. Those two lines intersect around λ = 0.35 1/h, defining the point of no drug. From this point, there are two ways to increase the ribosomal flux, *J_fr_* (the y-axis). You can either give the cells more food (leading to the black line, increasing *J_fr_* to the right) or give them drugs (leading to the blue line, increasing to the left). It suggests there are (at least) two signals that increase cellular ribosome fluxes: a signal about energy availability, and a signal about numbers of active ribosomes.

Related to that point, Fig. 6 shows the prediction of λ*_a_*(1–φ), which is a measure of the energy flux in the conversion of glucose to ATP, *m_a_k_a_* (*G*)ŀ*P*. Fig. 6 shows that there is a single universal relationship between that energy flux and growth rate, irrespective of whether growth is controlled by drugs or food. This indicates the nature of feedback in the cell. It is not simply the energy inflow (input) that dictates the growth rate (output). The growth rate is also a controller of the energy influx. This is interesting in the context of drugs, which can more strongly affect the growth rate dependence of the rate of ribosomal synthesis than energy influx. As far as we know, there are no experiments that bear on this prediction.

## 5. DISCUSSION

This model makes some predictions that have not yet been tested experimentally. We hope experimentalists will make such tests, to give deeper insights into these nonlinear behaviors, and to ultimately lead to improved models. Current experiments on drugged and undrugged bacteria are run on different food sources and in different media. Deeper tests of our model could come from studies that fix the types of nutrient and media, and vary only the food concentrations. In addition, a key variable here is *λ_a_*, the cell’s conversion efficiency of sugar to internal energy, such as ATP. It would be valuable to have measurements of: glucose and oxygen uptake rates, ATP production rates (*m_a_ J_a_*), ATP concentrations, and ribosome production rates (*J_r_*), key glycolytic, TCA cycle, and fermentation enzyme concentrations as a function of external glucose and antibiotic concentrations.

Somewhat different models are those of Elf et al. [35] and Deris et al. [36], who consider bistabilities of cells resulting either from membrane properties or drug resistance. Other models focus on mechanisms of microscopic control of ribosome synthesis, such as the “stringent response”, a negative feedback mechanism triggered when some of cell’s excess usable energetic molecules are converted to unusable ppGpp as response to endogenous limitations of aminoacids [21, 28, 37]. Because of its simplicity, the present treatment could be extended to explore other factors that are of interest, such as cellular geometry (surface-volume considerations), multi-drug effects [38], or drug-dependent cellular multistabilities that lead to antibiotic resistance and persistence [35, 36].

## 6. CONCLUSIONS

Here, we model the balance of energy, ribosomes and nonribosomal proteins in *E. coli* cells in the presence of chloramphenicol, an antibiotic drug. We suppose that chloramphenicol binds to ribosomes and inactivates them, in a Michaelis-Menten fashion. We combine this binding-induced inactivation of ribosomes with a three-component dynamical model of *E. coli’s* energy, ribosomal and non-ribosomal protein biomass as a function of growth rates, previously validated against experiments on undrugged bacteria. The present model gives quantitative predictions for how the cell’s growth rate decreases with drug, and how the total ribosomal fraction of protein increases with drug. And, it predicts that adding drugs to slow-growing cells leads to first increasing the rate of ribosomal synthesis, then a decrease as the cell gets sicker. We show the model agreement with data. But, more important are the insights the model gives about how the cell responds to drug: what varies and what stays constant. We find that while drugging the cell reduces the concentrations of active ribosomes, it also stimulates more total ribosome production, holding relatively constant the ribosomal production of ribosomes, a key quantity the cell uses to toggle between growth and self-protection.

## Author Contributions

A.M. and K.A.D designed research; A.M. performed research; A.M. and K.A.D analyzed data; and A.M. and K.A.D wrote the paper.

## Acknowledgement

We appreciate support from the Laufer Center at Stony Brook University.

## Supporting Information

### 1. RATE OF RIBOSOME CREATION

Here, we derive an expression for the rate of total ribosome production. Ribsomal inhibitors such as chloramphenicol reduce the cell’s growth rate by reducing the active fraction of ribosomes. Adding Eq. (2) and Eq. (13) and setting the sum to zero gives

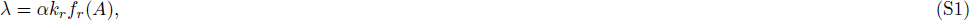

showing how the cell’s growth rate is reduced with *α*, the ribosomal inactivation fraction in Eq. 16.

Here, we express the rate of ribosome synthesis, *Jf_r_*, in units of g of ribosomal protein per g of total protein, to get the points on Fig. 5 as computed from experimental data *φ_tot_* – λ:

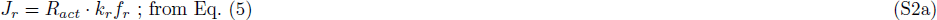

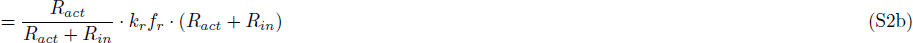

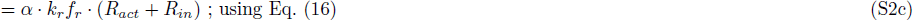

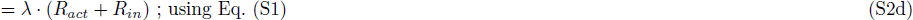

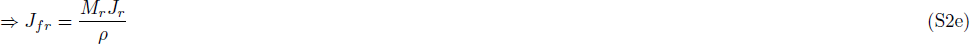

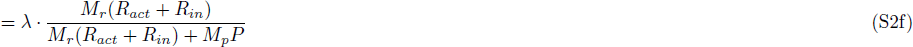

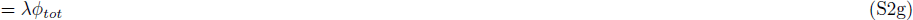

Under growth conditions without antibiotics, the flux for ribosomal synthesis is:

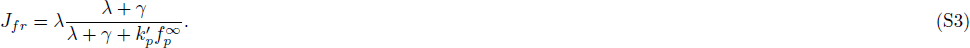

obtained from substituting Eq. (18) with α = 1, 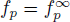 into Eq. (S2g).

### 2. ENERGY BALANCE

Next, we look at the energy balance. At steady state, Eqs. (1), (2), (3) are set to zero, leading to the expression,

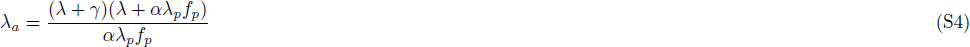

In addition, we now show how our model leads to the linear dependence of ribosomal content on growth rate described by Scott et al. [10]. We derive another expression for *φ_tot_* from Eqs. (18) and (S4) by eliminating 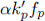:

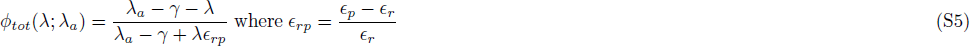

Setting 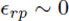 and 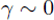 gives 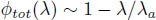.

**Table S1:** Structural, Rate and Bioenergetic Constants.

| Physical constants | Symbol | Value |
| --- | --- | --- |
| Protein density | $\rho$ | 0.25 g cm<sup>-3</sup> |
| Molec. wt. of ribosomal proteins (RP) per ribosome | $M_r$ | 7336 aa × 110 g/mol/aa = 806960 g mol<sup>-1</sup> |
| Molec. wt. of a non-ribosomal protein (NRP) | $M_p$ | 325 aa × 110 g/mol/aa = 35750 g mol<sup>-1</sup> |
| Molecules of ATP produced per glucose molecule | $m_a$ | 30 |
| Molecules ATP consumed to create one ribosome | $m_r$ | (7336 aa × 6) + (4566 nu × 10) ∼ 89700 |
| Molecules of ATP consumed to create one NRP | $m_p$ | 325 aa × 6 = 1950 |
| Rate of NRP elongation per ribosome, 20 aa/s | $k'_p$ | 20 × 3600 (aa/h)/7336 aa ∼ 10 aa/h/(RP aa) |
| Non-ribosomal protein degradation rate | $\gamma$ | 0.1 NRP per total NRP per h |

| Derived constants | Symbol | Value |
| --- | --- | --- |
| Max no. of protein molecules translated per hr per ribosome (capacity) | $k_p$ | $M_r k'_p / M_p = 215 \text{ h}^{-1}$ |
| NRP translation rate per ribosome scaled by pathway efficiencies | $\lambda_p$ | $(\varepsilon_r / \varepsilon_p) k'_p \sim 5 \text{ h}^{-1}$ |
| Max no. of ribosomes synthesized per hr per ribosome ($= \lambda_p$) | $k_r$ | 5 h<sup>-1</sup> |
| Ribosomal pathway efficiency, g of RPs synthesized per mol ATP | $\varepsilon_r$ | $M_r / m_r \sim 9 \text{ g mol}^{-1}$ |
| Protein pathway efficiency, g of NRPs per mol ATP | $\varepsilon_p$ | $M_p / m_p \sim 18 \text{ g mol}^{-1}$ |
| Relative pathway efficiency between P- and R- pathways | $\varepsilon_{rp}$ | $(\varepsilon_p - \varepsilon_r) / \varepsilon_r \sim 1$ |

**Table S2:** Parameters of *E. coli* ODE numerical model obtained from fit of the model to data.

| ODE model Parameters | Symbol | Value |
| --- | --- | --- |
| Affinity constant between nonribosomal proteins and glucose for glucose transport | $D_g$ | 0.07 mM |
| Number of glucose molecules metabolized to ATP per hr per protein molecule | $k_a^\infty$ | 120 h<sup>-1</sup> |
| Affinity constant between proteins and ATP for ATP generation | $D_a$ | 4.0 mM |
| ATP concentration threshold for ribosome synthesis | $D_r$ | 0.18 mM |
| Max fraction of ribosomes translating RPs | $f_r^\infty$ | 0.2 |
| Max fraction of ribosomes translating NRPs | $f_p^\infty$ | 0.7 |

**Table S3:**
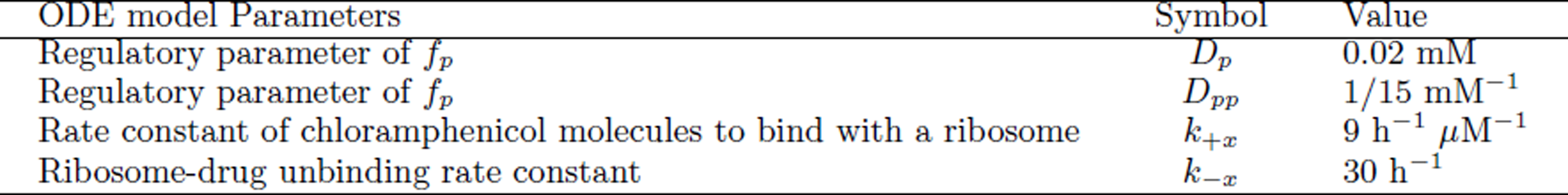
Interaction parameters of E. coli and Chloramphenicol.

| ODE model Parameters | Symbol | Value |
| --- | --- | --- |
| Regulatory parameter of $f_p$ | $D_p$ | 0.02 mM |
| Regulatory parameter of $f_p$ | $D_{pp}$ | 1/15 mM<sup>-1</sup> |
| Rate constant of chloramphenicol molecules to bind with a ribosome | $k_{+x}$ | 9 h<sup>-1</sup> $\mu\text{M}^{-1}$ |
| Ribosome-drug unbinding rate constant | $k_{-x}$ | 30 h<sup>-1</sup> |

**Figure S1:**
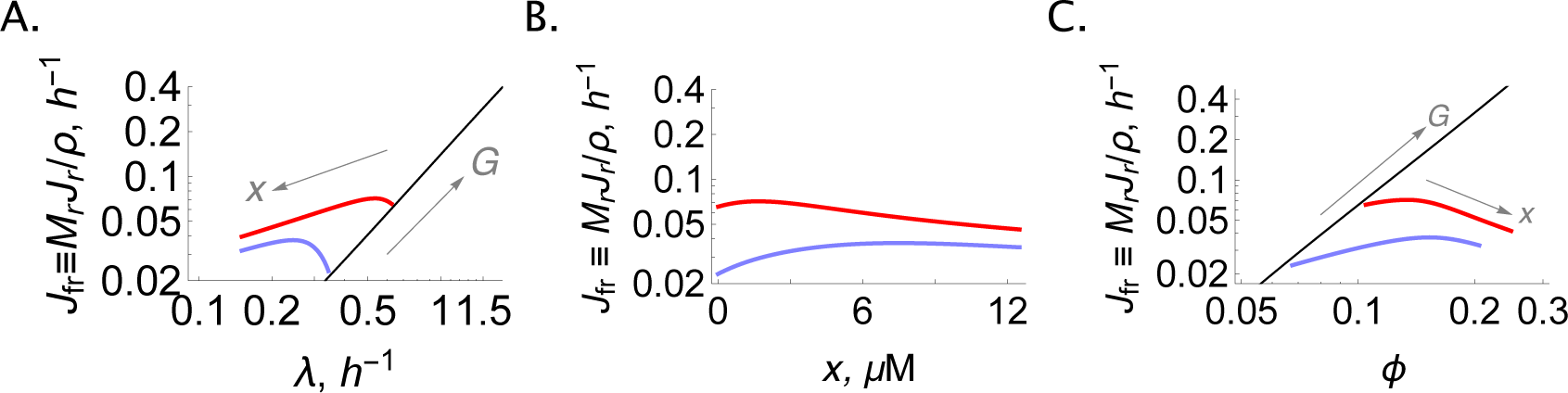
Effect of ribosomal inhibitors on ribosomal activity. (A) The black line is theory, Eq. (S3), in the absence of drugs with 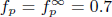, 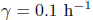, 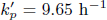. Solutions of ODE cell model at glucose concentrations *G*=0.04 mM (blue) and 0.08 mM (red), respectively, under increasing chloramphenicol concentrations, *x* = 0 – 15 *μ*M, shown by arrow. (B) Ribosomal activity vs. chloramphenicol concentration from ODE model at *G*=0.04 mM (blue) and 0.08 mM (red). (C) Ribosomal activity vs. ribosomal protein fraction from ODE model at *G*=0.04 mM (blue) and 0.08 mM (red).

**Figure S2:**
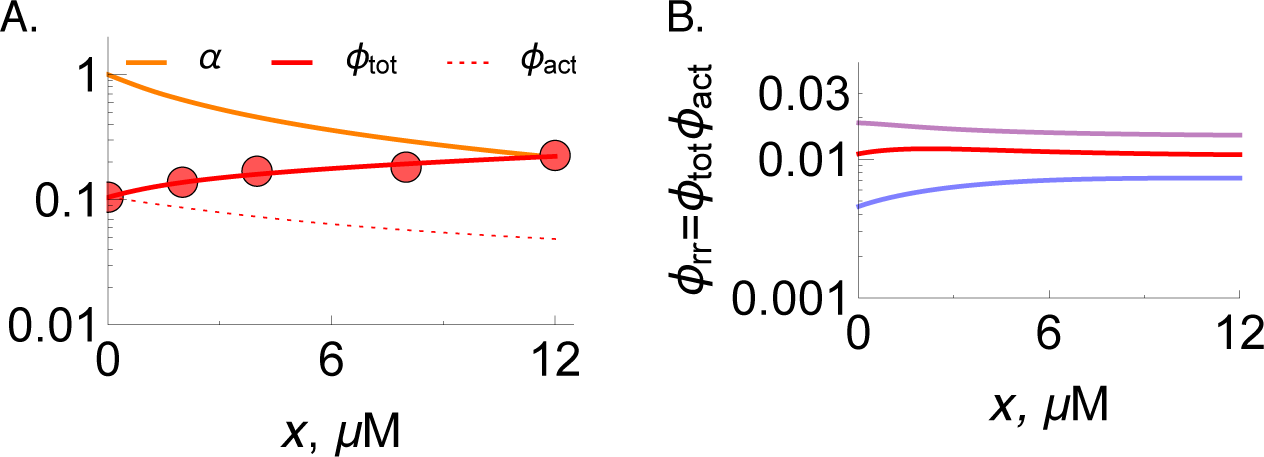
Effect of ribosomal inhibitors on cellular homeostasis. (A) Solid circles are experimental data of ribosomal protein fraction *φ_tot_* of *E. coli* grown on glucose+M63 [10, 32]. Lines are numerical solutions of the ODE model, with glucose concentration *G* = 0.08 mM. Orange, *α* active fraction of ribosomes independent of *G* and dependent hyperbolically on chloramphenicol concentration *x*; see Eq. (16). Red solid line, total ribosomal protein fraction, 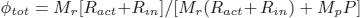, at different dosage of chloramphenicol, *x*. Red dotted line, 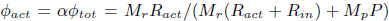 Eq. (17). (B) 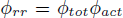, fraction of active ribosomes devoted to ribosomal autosynthesis, from model for *G* = 0.04 mM (blue), 0.08 mM (red), 0.125 mM (purple).

**Figure S3:**
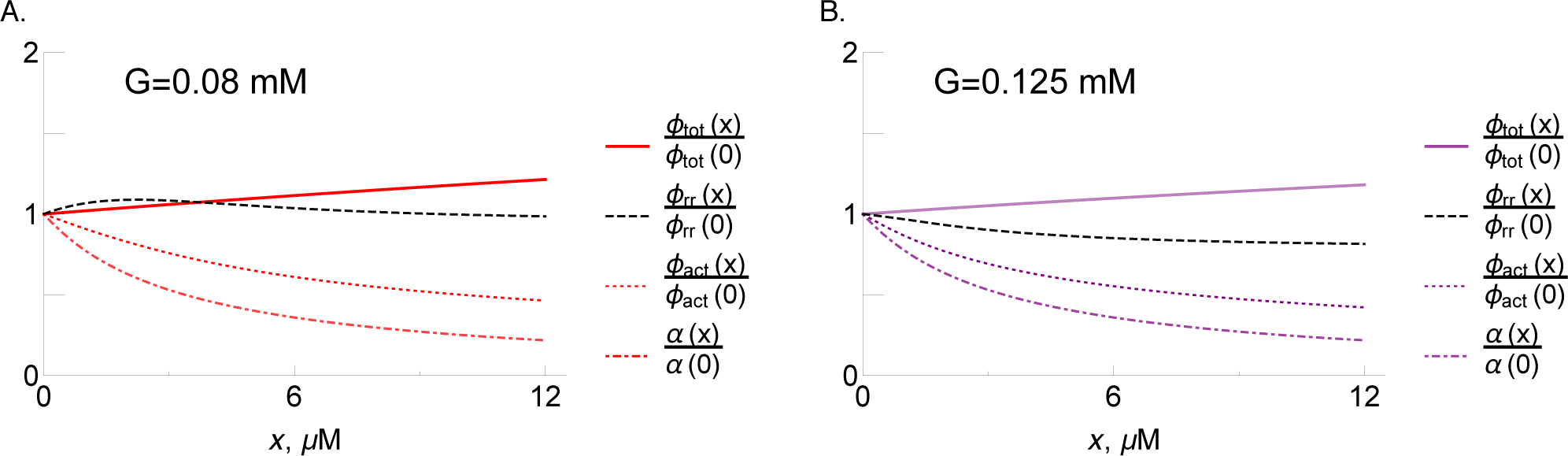
Effect of ribosomal inhibitors on cellular homeostasis. Numerical solutions of the ODE model under varying concentrations of ribosomal inhibitors *x* and fixed glucose concentrations: (A) *G* = 0.08 mM, (B) 0.125 mM. Solid lines, total ribosomal protein fraction, 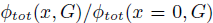. Black dashed lines, 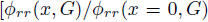. Dotted lines, 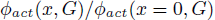; from Eq. (17). Dot-dashed lines, *α*, active fraction of ribosomes; see Eq. (16).

**Figure S4:**
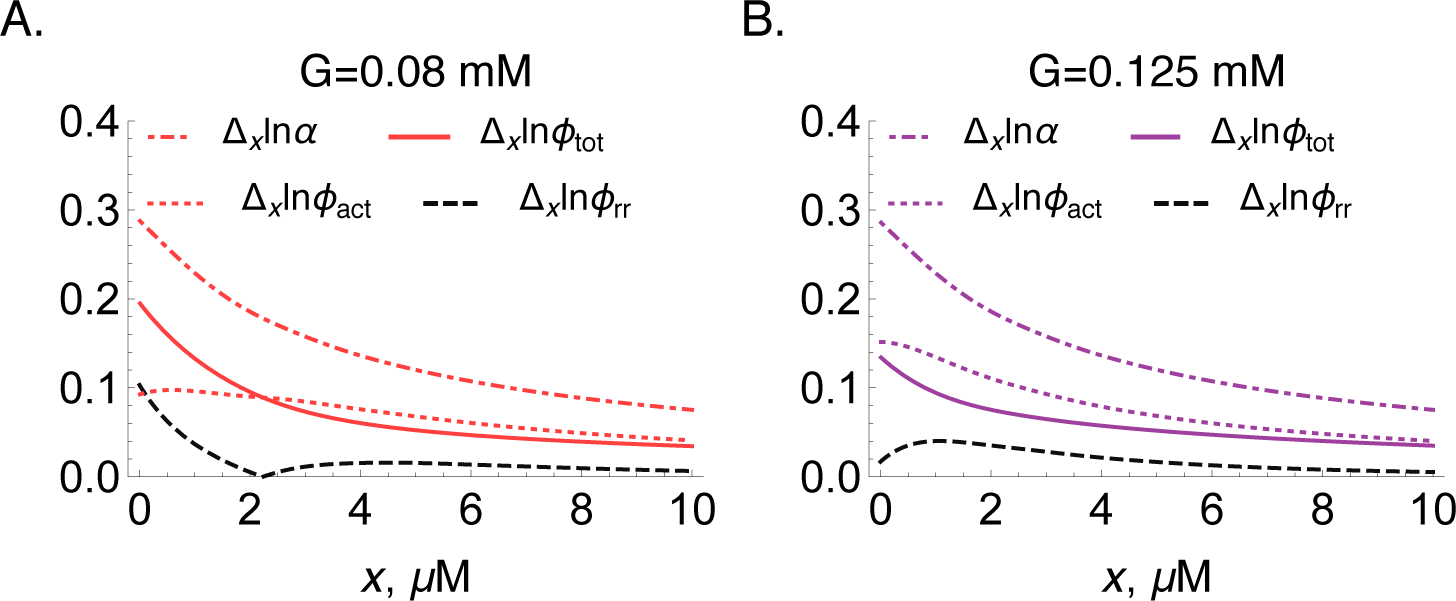
Relative change in ribosomal protein fractions with change in inhibitor concentrations. A comparison of the quantities 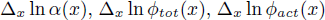 and 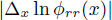 where 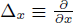 obtained from model at fixed glucose concentrations: (A) *G* = 0.08 mM, and (B) *G* = 0.125 mM. These plots show that the quantity 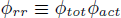 is more tightly regulated compared to the other quantities under antibiotic stress.

